# GDF15 Is Required for Cold-Induced Thermogenesis and Contributes to Improve Systemic Metabolic Health Following Loss of OPA1 in Brown Adipocytes

**DOI:** 10.1101/2022.12.23.521796

**Authors:** Jayashree Jena, Luis Miguel García-Peña, Eric T. Weatherford, Alex Marti, Sarah H. Bjorkman, Kevin Kato, Jivan Koneru, Jason Chen, Randy J. Seeley, E. Dale Abel, Renata O. Pereira

## Abstract

Previously we showed that mice lacking the protein optic atrophy 1 (OPA1 BKO) in brown adipose tissue (BAT) have induction of the activating transcription factor 4 (ATF4), which promotes fibroblast growth factor 21 (FGF21) secretion as a batokine. FGF21 increases metabolic rates at baseline conditions but is dispensable for the resistance to diet-induced obesity (DIO) reported in OPA1 BKO mice [1]. To determine alternative mediators of this phenotype, we performed transcriptome analysis, which revealed increased levels of growth differentiation factor 15 (GDF15) in BAT. To determine if ATF4 induction was mediated by the protein kinase R (PKR)-like endoplasmic reticulum kinase (PERK), and to evaluate the contribution of GDF15 to the resistance to DIO, we selectively deleted PERK or GDF15 in OPA1 BKO mice. Mice lacking both OPA1 and PERK in BAT had preserved induction of ATF4. Importantly, simultaneous deletion of OPA1 and GDF15 partially reversed the resistance to diet-induced obesity and abrogated the improvements in glucose tolerance. Furthermore, GDF15 mediated cold-induced thermogenesis, likely via sympathetic activation of iWAT. Taken together, our data indicate that PERK is dispensable for ATF4 induction, but GDF15 contributes to the resistance to DIO, and is required for glucose homeostasis and thermoregulation in OPA1 BKO mice.

## Introduction

The integrated stress response (ISR) is a pro-survival signaling pathway present in eukaryotic cells, which is activated in response to a range of physiological and pathological stressors. Such stresses commonly include cell extrinsic factors such as hypoxia, amino acid deprivation, glucose deprivation, and viral infection. However, cell intrinsic stresses such as endoplasmic reticulum (ER) stress [2], and mitochondrial stress can also activate the ISR [1, 3, 4]. Induction of the ISR and its main effector activating transcription factor 4 (ATF4) in response to different stress conditions frequently correlates with increased levels of the endocrine factors fibroblast growth factors 21 (FGF21) and growth differentiation factor 15 (GDF15) in various tissues, including in liver [5] and in skeletal muscle [6]. However, the upstream mechanisms mediating ISR induction in response to mitochondrial stress are unclear. Furthermore, the respective roles of FGF21 and GDF15 in the context of mitochondrial dysfunction are incompletely understood.

We recently reported that mitochondrial stress caused by deletion of the mitochondrial fusion protein optic atrophy 1 (OPA1 BAT KO) activated the ISR and induced its main effector the activating transcription factor 4 (ATF4) in brown adipose tissue (BAT). ATF4-mediated FGF21 induction was required to promote browning of inguinal white adipose tissue (iWAT), and to improve thermoregulation at baseline conditions. Although activation of the ISR in BAT correlated with resistance to diet-induced obesity (DIO) in an ATF4-dependent manner, this phenomenon was independent of FGF21 [1]. In the present study, we sought to investigate the upstream mechanisms mediating ISR activation in BAT in response to OPA1 deletion, and the molecular mechanisms downstream of ATF4 mediating resistance to DIO.

Transcriptome analysis showed that the unfolded protein response (UPR) was induced in OPA1-deficient BAT. Given that the protein kinase R (PKR)-like ER kinase (PERK) is shared by the ISR and the UPR [2], here, we tested the hypothesis that PERK is required for ATF4 induction in BAT. We also observed that in addition to FGF21, GDF15 was highly induced in BAT upon OPA1 deletion. Because GDF15 has been shown to regulate energy homeostasis in rodents [7], here we tested the hypothesis that GDF15 is downstream of ATF4 and mediates the resistance to DIO in mice lacking OPA1 in thermogenic adipocytes. Our data reveal that PERK is dispensable for ISR and ATF4 induction in BAT in response to OPA1 deletion. Notably, GDF15 deficiency does not prevent the induction of browning, but is required for thermogenic activation of iWAT likely by mediating its sympathetic activation. Despite this, GDF15 only partially mediates the resistance to DIO, although it is necessary for the improvement in glucose homeostasis and hepatic steatosis following high-fat feeding in mice deficient for OPA1 in BAT. These data underscore the complex regulation of systemic metabolism by various BAT-derived endocrine regulators that are released in response to mitochondrial stress.

## Results

### Transcriptome Analysis Reveals Induction of the UPR and GDF15 in Mice Lacking OPA1 in BAT

To gain insight into the molecular mechanisms regulating activation of the ISR in BAT in response to OPA1 deletion, and the mechanisms downstream of ATF4 that mediate resistance to DIO in mice lacking OPA1 in BAT, we performed RNA sequencing (RNASeq) in BAT collected from 7-week-old male and female mice lacking OPA1 in thermogenic adipocytes (OPA1 BAT KO) and their respective wild-type (WT) littermate controls at ambient temperature conditions. Ingenuity Pathway Analysis revealed that the ER stress pathway and the UPR were amongst the top-3 upregulated canonical pathways in OPA1-deficient BAT (Fig. 1A).

**Figure 1.**
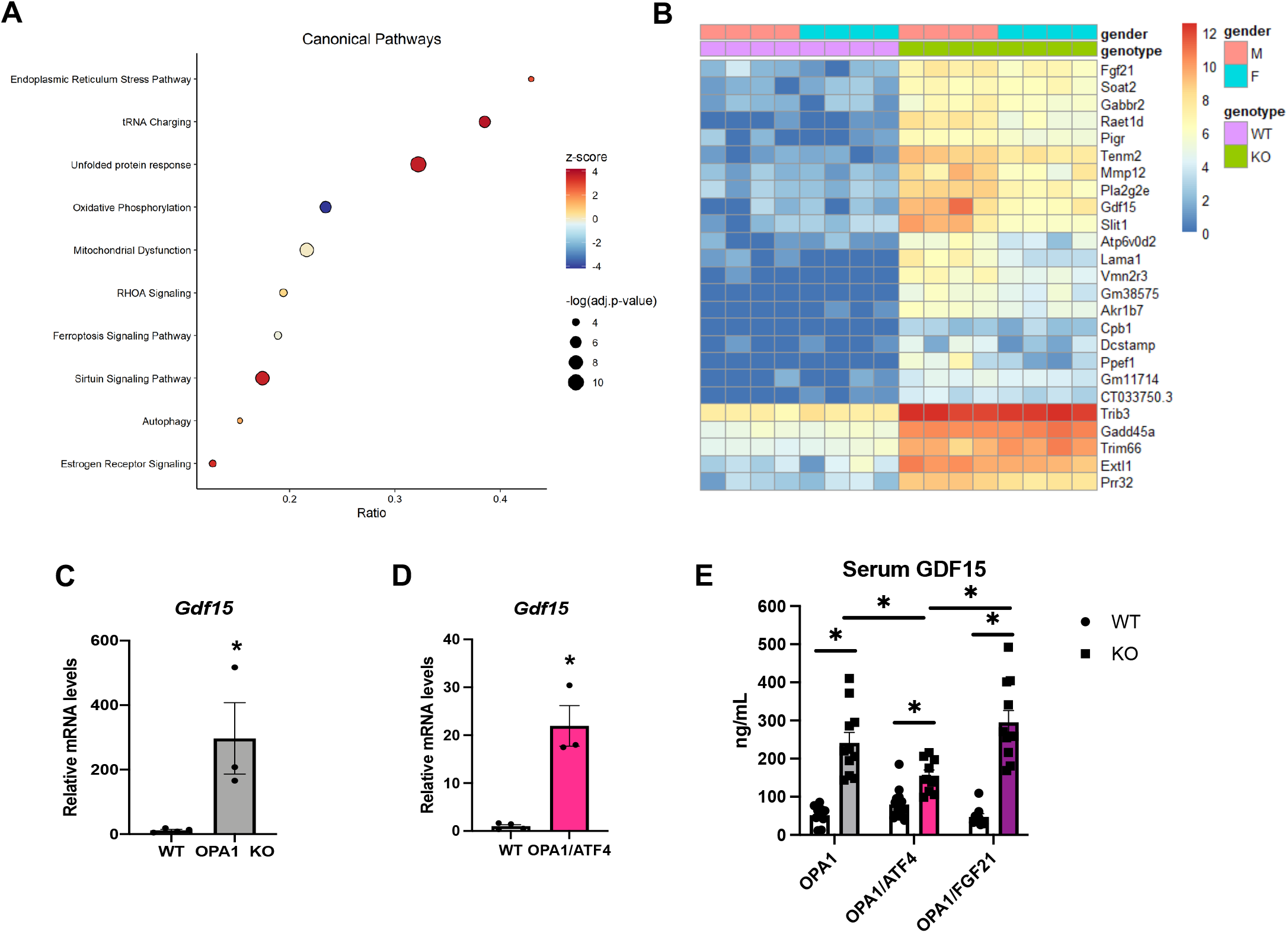
RNASeq analysis and Gdf15 expression in OPA1 BAT KO mice: A-D. Data collected from brown adipose tissue (BAT) of 7-week-old male and female OPA1 BAT KO mice. **A**. Bubble plot showing the top 10 canonical pathways from the IPA database containing genes with a significant overlap (adjusted p-value ≤ 0.05) to those differentially expressed in OPA1 BAT KO mice. Size of the bubble indicates the -log of the adjusted p-value (Benjamini-Hochberg) from the pathway analysis. Plotted on the x-axis is the ratio of differentially expressed genes relative to the number of genes in the pathway. The bubble color indicates the z-score which indicates the predicted activation (positive) or inhibition (negative) of the pathway based on the directionality of the gene changes in OPA1 BAT KO relative to wild type (WT) mice. **B**. Heatmap of the top 25 differentially expressed genes in OPA1 BAT KO mice. **C**. Relative mRNA expression of *Gdf15* in BAT of OPA1 BAT KO mice normalized to *Gapdh* expression. **D**. Relative mRNA expression of *Gdf15* in BAT of OPA1/ATF4 BAT DKO mice normalized to *Gapdh* expression. **E**. GDF15 serum levels in OPA1 BAT KO, OPA1/ATF4 BAT DKO and OPA1/FGF21 BAT DKO mice. Data are expressed as means ± SEM. Significant differences were determined by Student’s *t*-test, using a significance level of p<0.05. *Significantly different vs. WT mice.

Furthermore, in addition to FGF21, which we previously demonstrated was required for activation of browning of iWAT and to regulate changes in core body temperature at baseline conditions, but not for the resistance to DIO in OPA1 BAT KO mice [1], GDF15 was amongst the top 25 up-regulated genes in BAT in response to OPA1 deletion (Fig. 1B). *Gdf15* mRNA induction in OPA1 BAT KO mice was confirmed by qPCR (Fig. 1C). Interestingly, *Gdf15* mRNA levels were still significantly induced in mice lacking both ATF4 and OPA1 in BAT (Fig. 1D), although the fold-change increase relative to WT mice was ∼ 100 times reduced compared to mice with OPA1 deletion alone (Fig. 1C). Importantly, GDF15 serum levels were induced in OPA1 BAT KO mice but were significantly attenuated in mice lacking both OPA1 and ATF4 in BAT (OPA1/ATF4) (Fig. 1E). Conversely, GDF15 serum levels remained elevated in mice concomitantly lacking OPA1 and FGF21 in BAT (Fig. 1E). These are important control groups, as mice lacking both OPA1 and ATF4 in BAT are no longer resistant to DIO, while mice lacking OPA1 and FGF21 in BAT remain lean when fed high-fat diet (HFD) [1].

### PERK Is Dispensable for the ISR Activation in OPA1 BAT KO Mice

Because the ER kinase PERK is shared by the UPR and the ISR and leads to induction of ATF4 [2], we hypothesized that activation of the PERK arm of the UPR is required to activate the ISR in OPA1 BAT KO mice. To test this hypothesis, we generated mice concomitantly lacking both OPA1 and PERK in thermogenic adipocytes (OPA1/PERK BAT DKO) (Fig. 2A). In the absence of PERK, phosphorylation of the translation initiation factor eIF2α, which is required for ATF4 translation, remained elevated in the OPA1 BAT KO background (Fig. 2B). Accordingly, induction of the ISR genes *Atf4, Ddit3, Fgf21* and *Gdf15* remained elevated in BAT (Fig. 2C), which correlated with a significant increase in GDF15 serum levels (Fig. 2D). Metabolically, OPA1/PERK BAT DKO mice, as was also the case with OPA1 BAT KO mice, had reduced body mass (Fig. 2E) and total fat mass (Fig. 2F), with unchanged total lean mass (Fig. 2G) at 20 weeks of age. Uncoupling protein 1 (UCP1) levels in the inguinal white adipose tissue (iWAT) were also elevated in OPA1/PERK BAT DKO, indicating increased baseline browning of iWAT (Fig. 2H). After 12 weeks of high-fat feeding, OPA1/PERK BAT DKO mice were resistant to DIO, as demonstrated by reduced body mass (Fig. 2I) and total fat mass (Fig. 2J). ANCOVA analysis revealed a leftward shift in the relationship of energy expenditure as a function of body mass in the OPA1/PERK BAT DKO mice, with a significant difference in the group effect (Fig. 2K). These data indicate increased energy expenditure in OPA1/PERK BAT DKO mice. Glucose (Fig. 2L and M) and insulin intolerance (Fig. 2N and O) were ameliorated in high-fat-fed OPA1/PERK BAT DKO, as was also the case in OPA1 BAT KO mice. Thus, PERK is dispensable for the ISR activation in BAT in response to OPA1 deletion, and for the associated systemic metabolic improvements. Therefore, alternative ISR kinases are required to activate the ISR in response to mitochondrial stress.

**Figure 2.**
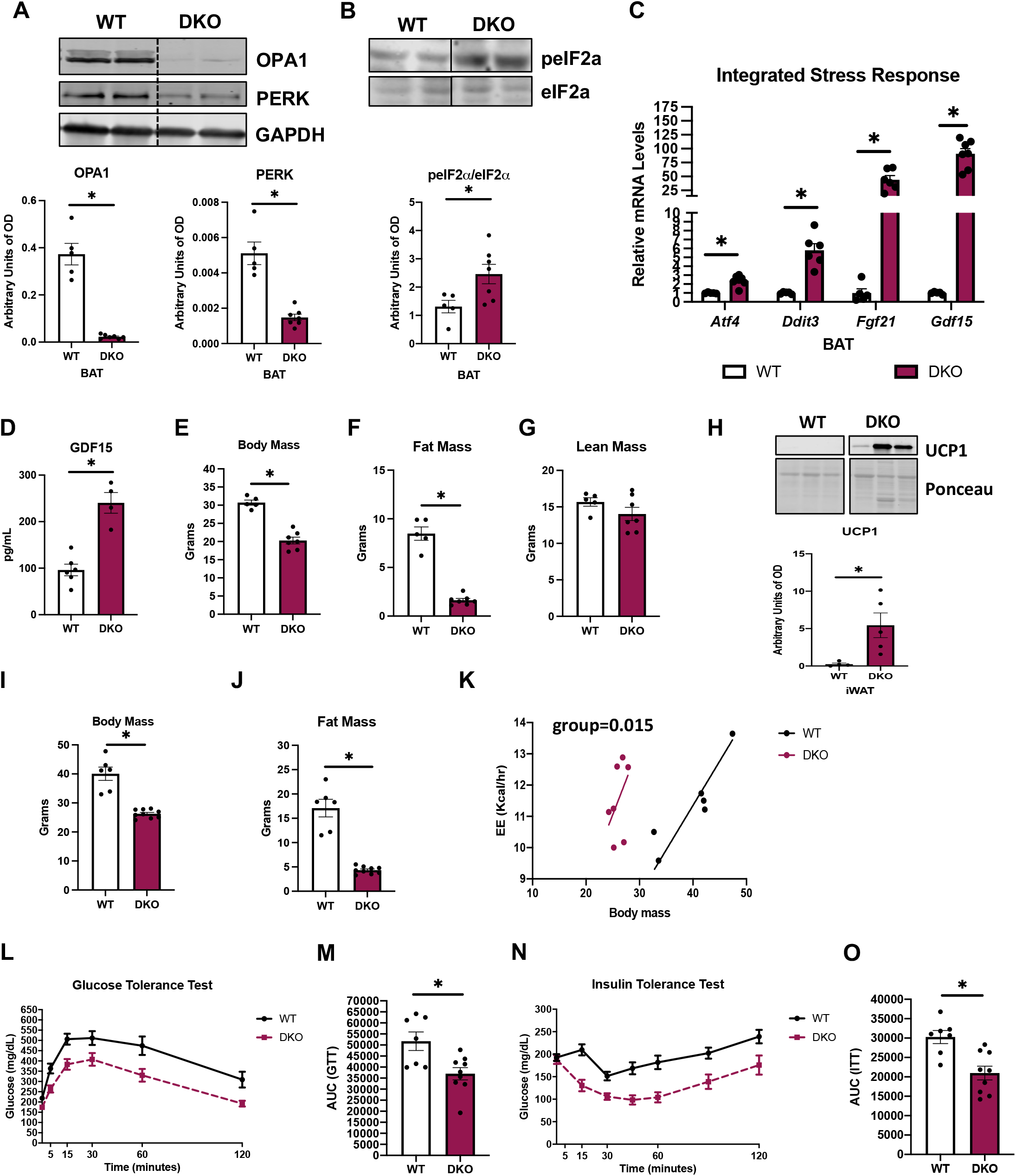
PERK Is Dispensable for the ISR Activation in OPA1 BAT KO Mice: A-D. Data collected from 8-week-old OPA1/PERK BAT double-knockout (DKO) mice. **A**. Representative immunoblots for OPA1 and PERK protein in BAT normalized to GAPDH, and their respective densitometric quantification. **B**. Representative immunoblots for phospho-eIF2α (peIF2α) normalized to total eIF2α protein levels in BAT and the respective densitometric quantification. Optical density (OD). **C**. Relative mRNA expression of the integrated stress response (ISR) genes *Atf4, Ddit3, Fgf21* and *Gdf15* normalized to *Tbp* expression in BAT. **D**. GDF15 serum levels (*ad libitum*-fed conditions). **E-H**. Data collected in 20-week-old DKO mice. **E**. Body mass. **F**. Total fat mass. **G**. Total lean mass. **H**. Representative immunoblots for the uncoupling protein 1 (UCP1) normalized to ponceau staining, and the respective densitometric quantification. **I-O**. Data collected in DKO mice fed a high-fat diet (HFD) for 12 weeks. **I**. Body mass. **J**. Total fat mass. **K**. Regression plot of energy expenditure (EE) as a function of body mass. **L**. Glucose tolerance test (GTT). **M**. Area under the curve (AUC) for the GTT. **N**. Insulin tolerance test (ITT). **O**. Area under the curve (AUC) for the ITT. Data are expressed as means ± SEM. Significant differences were determined by Student’s *t*-test, using a significance level of p<0.05. Energy expenditure data was analyzed by ANCOVA. *Significantly different vs. WT mice.

### GDF15 Partially Mediates the Resistance to DIO, and Is required to Improve Glucose Homeostasis in OPA1 BAT KO Mice

Our transcriptome data show that *Gdf15* mRNA levels are highly induced in OPA1 BAT KO mice, and its induction is attenuated in the absence of ATF4, suggesting partial dependence on ATF4 for GDF15 regulation (Fig. 1F and G). Given GDF15’s known effects on energy metabolism in rodents [8], we tested the hypothesis that GDF15 induction in OPA1 BAT KO is required to mediate the resistance to DIO. For that, we generated mice with simultaneous deletion of both the *Opa1* and *Gdf15* genes in thermogenic adipocytes (OPA1/GDF15 BAT DKO). mRNA expression demonstrated a significant reduction in *Opa1* and *Gdf15* levels in BAT of DKO mice (Fig. 3A), which completely normalized GDF15 serum levels (Fig. 3B). mRNA expression of the ISR genes *Atf4, Ddit3* and *Fgf21* remained elevated in BAT of DKO mice (Fig. 3C), which correlated with increased FGF21 serum levels (Fig. 3D). At baseline conditions, body mass was unchanged at 6 and 10 weeks of age but was significantly reduced in 20-week-old DKO mice, relative to controls (Fig. 3E). At 10-12 weeks, although body mass was unchanged, energy expenditure was slightly increased in DKO mice during the dark cycle (Fig. 3F), while food intake (Fig. 3G), and locomotor activity (Fig. 3H) were unchanged. These data suggest that GDF15 does not regulate changes in energy expenditure in OPA1 BAT KO mice at baseline conditions, when mice were fed isocaloric diet. Indeed, our published data suggest FGF21 regulates changes in energy homeostasis and body mass in OPA1 BAT KO mice under these conditions [1]. We next tested whether BAT-derived GDF15 is required to promote resistance to DIO in OPA1 BAT KO mice. WT and DKO mice were fed a HFD (60% calories from fat) for 12 weeks. Body mass (Fig. 3I) and total fat mass (Fig. 3J) were significantly reduced in DKO mice, while lean mass was unchanged (Fig. 3K). Although body mass was significantly reduced in DKO mice, when compared to the final body mass of OPA1 BAT KO subjected to the same DIO protocol (data previously published by us), DKO mice gained significantly more weight (Fig. 3L). These data suggest that GDF15 partially mediates the resistance to DIO observed in OPA1 BAT KO mice. To gain insight into the mechanisms contributing to these changes in weight gain, we placed a subset of high-fat fed WT and DKO in metabolic chambers. Surprisingly, energy expenditure (Fig. 3M), food intake (Fig. 3N) and locomotor activity (Fig. 3O) were all unchanged between genotypes. Indeed, the group effect from the ANCOVA analysis for oxygen consumption rates in relation to body weight was not significantly different (Fig. 3P), suggesting GDF15 may mediate the increase in metabolic rates observed in OPA1 BAT KO mice in DIO. Although body mass was significantly reduced in DKO mice relative to WT mice, glucose homeostasis, as demonstrated by the glucose tolerance test (Fig. 3Q and R) and fasting glucose levels (Fig. 3S) were similarly impaired in WT and DKO mice. Likewise, diet-induced hepatic triglyceride accumulation was comparable between WT and DKO (Fig. 3T). Conversely, insulin sensitivity was significantly improved in DKO mice, as shown by the reduced area under the curve for the insulin tolerance test (Fig. 3U and V) and decreased serum fasting insulin levels (Fig. 3W).

**Figure 3.**
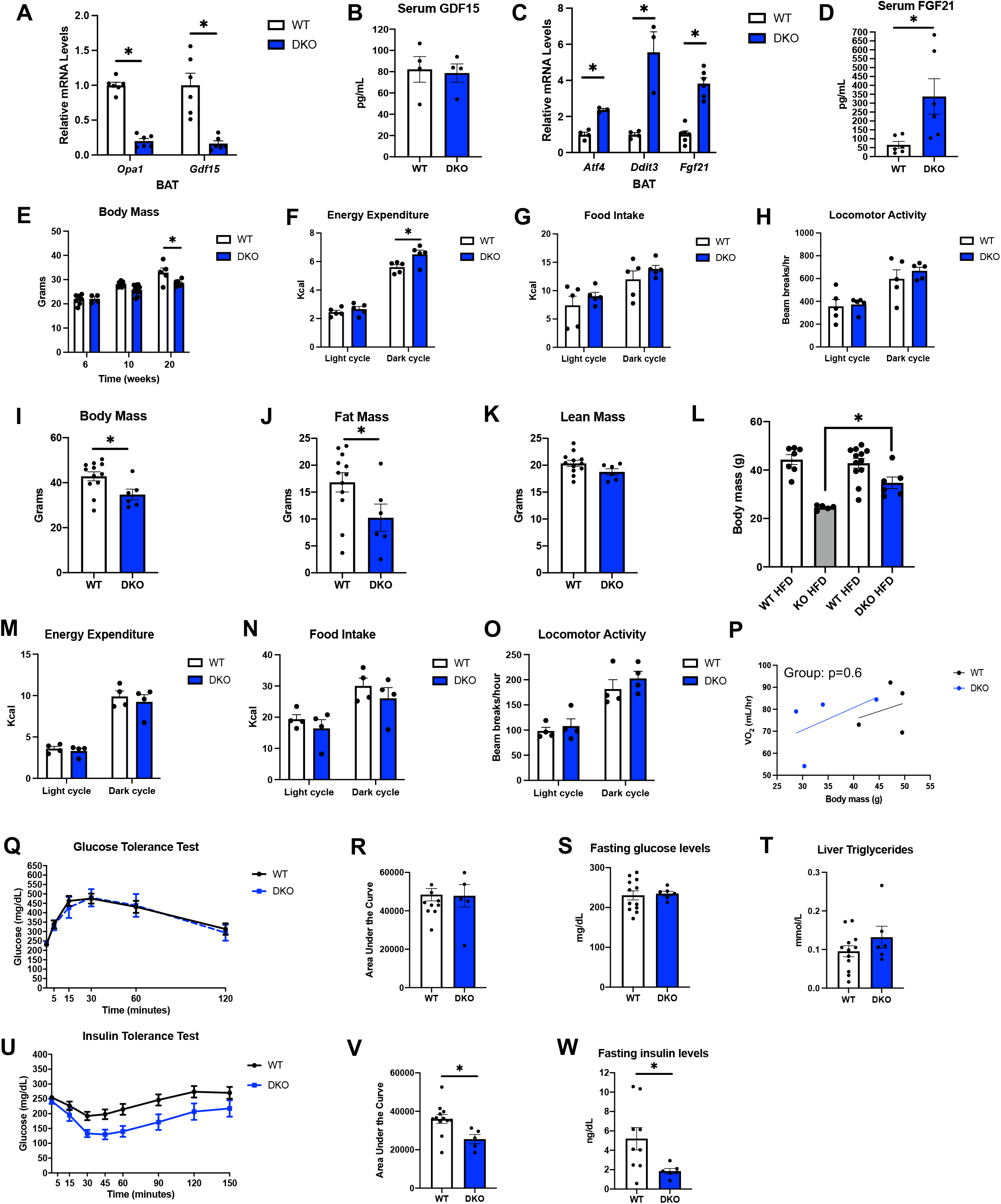
GDF15 Partially Mediates the Resistance to DIO and Is required to Improve Glucose Homeostasis in OPA1 BAT KO Mice. A-H. Data collected from OPA1/GDF15 BAT double-knockout (DKO) mice fed isocaloric diet. **A**. Relative mRNA expression of *Opa1* and *Gdf15* in BAT normalized to *Tbp* expression. **B**. GDF15 serum levels (*ad libitum*-fed conditions). **C**. Relative mRNA expression of the integrated stress response (ISR) genes *Atf4, Ddit3* and *Fgf21* normalized to *Tbp* expression. **D**. FGF21 serum levels (*ad libitum*-fed conditions). **E**. Body mass at 8, 10 and 20 weeks of age. **F**. Energy expenditure in 10-12-week-old mice (before changes in body mass were detected). **G**. Food intake. **H**. Locomotor activity. **I-W**. Data collected in DKO mice fed a high-fat diet (HFD) for 12 weeks. **I**. Body mass. **J**. Total fat mass. **K**. Total lean mass. **L**. Direct comparison between body mass in OPA1 BAT KO mice (KO) and OPA1/GDF15 BAT DKO mice (DKO). **M**. Energy expenditure. **N**. Food intake. **O**. Locomotor activity. **P**. Regression plot of oxygen consumption (VO_2_) as a function of body mass. **Q**. Glucose tolerance test (GTT). **R**. Area under the curve (AUC) for the GTT. **S**. Fasting glucose levels (collected after 6-hr fast). **T**. Liver triglyceride levels. **U**. Insulin tolerance test (ITT). **V**. Area under the curve (AUC) for the ITT. **W**. Fasting insulin levels. Data are expressed as means ± SEM. Significant differences between two groups were determined by Student’s *t*-test, using a significance level of p<0.05. Significant differences between 3 or more groups were determined by two-Way ANOVA using a significance level of p<0.05. *Significantly different vs. WT mice, or significantly different from KO HFD. VO_2_ data was analyzed by ANCOVA.

### GDF15 Is Required to Regulate Core Body Temperature in Cold-Exposed OPA1 BAT KO Mice

Baseline expression of thermogenic genes was reduced in the BAT of DKO mice (Fig. 4A), suggesting BAT thermogenesis is impaired in these mice, as is the case in OPA1 BAT KO mice. We, therefore, tested whether core body temperature would be affected in DKO mice at thermoneutrality and in response to cold stress. After 7 days at 30 °C, core body temperature was slightly increased in DKO mice relative to their WT littermate controls (Fig. 4B). Conversely, when housed at 4 °C, core body temperature, as measured by telemetry, was reduced in DKO mice relative to their WT counterparts (Fig. 4C). These changes in temperature occurred independently of changes in energy expenditure (Fig. 4D) or food intake (Fig. 4E) between genotypes (mice that died within the first 24 hours of cold exposure were removed from the analysis). To capture the changes in core body temperature in all mice, we also plotted the last temperature recorded per mouse, which was significantly reduced in DKO mice (Fig. 4F). In a separate cohort of mice, we used a rectal probe to measure core body temperatures prior to cold exposure (time 0) and hourly after ambient temperatures reached 4 °C (for up to 5 hours).

**Figure 4.**
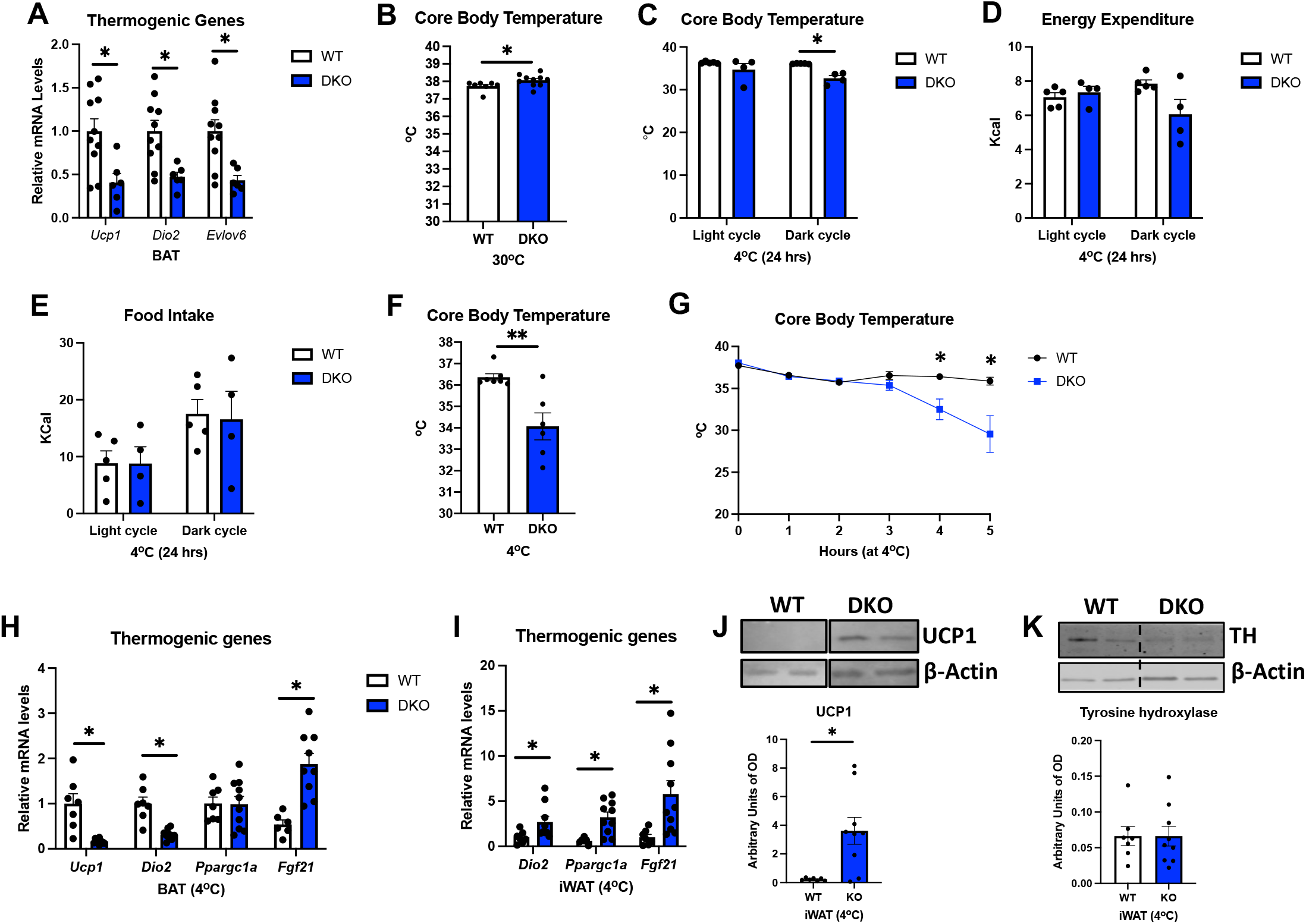
GDF15 Is Required to Regulate Core Body Temperature in Cold-Exposed OPA1 BAT KO Mice. **A**. mRNA expression of the thermogenic genes *Ucp1, Dio2* and *Evlov6* in BAT of OPA1/GDF15 BAT DKO mice (DKO) at ambient temperature conditions. **B**. Core body temperatures collected from 12-week-old wild type (WT) and DKO mice after 7 days at 30 °C. **C**. Averaged core body temperature (light and dark cycles) in mice cold exposed for 24 hours (4 °C). **D**. Averaged energy expenditure (light and dark cycles) in mice, cold exposed for 24 hours (4 °C). **E**. Averaged food intake (light and dark cycles) in mice, cold exposed for 24 hours (4 °C). **F**. Final core body temperature recorded by telemetry in mice exposed to 4°C in the CLAMS system. **G**. Hourly core body temperatures collected from 12-week-old wild type (WT) and DKO mice after cold exposure (4°C). **H**. mRNA expression of thermogenic genes in BAT after 5 hours of cold exposure. **I**. mRNA expression of thermogenic genes in inguinal white adipose tissue (iWAT) after 5 hours of cold exposure. **J**. Representative immunoblots for UCP1 in iWAT after 5 hours of cold exposure normalized to β-actin, and their respective densitometric quantification. **K**. Representative immunoblots for tyrosine hydroxylase (TH) in iWAT of mice cold-exposed for 5 hours normalized to β-actin, and their respective densitometric quantification. Optical density (OD). Data are expressed as means ± SEM. Significant differences were determined by Student’s *t*-test, using a significance level of p<0.05. *Significantly different vs. WT mice.

Our data shows a precipitous decline in core body temperature in DKO mice, starting at 4 hours post-cold exposure (Fig. 4G). After 5 hrs., animals were euthanized, and tissues were harvested for analysis. mRNA levels of *Ucp1* and *Dio2* were significantly reduced in BAT of DKO mice, while *Fgf21* mRNA levels remained elevated (Fig. 4H). Surprisingly, mRNA expression of thermogenic genes (Fig. 4I) and protein levels of UCP1 were induced in the inguinal white adipose tissue (iWAT) of DKO mice (Fig. 4J), suggesting increased browning. However, protein levels of tyrosine hydroxylase in iWAT, a proxy for sympathetic activation, were unchanged between WT and DKO mice (Fig. 4K).

These data suggest that, although GDF15 induction in BAT is dispensable for the baseline increase in core body temperature observed in OPA1 BAT KO mice, it is required to regulate core body temperature in cold-exposed OPA1 BAT KO mice. The mechanism likely reflects a role for GDF15 in regulating sympathetic activation of iWAT and suggest that induction of thermogenic genes alone without the accompanying increase in sympathetic innervation is insufficient to sustain thermogenesis in mice lacking OPA1 in thermogenic adipocytes.

## Discussion

In our recently published study, we showed that deletion of the mitochondrial fusion protein OPA1 in brown adipocytes impaired thermogenic capacity in BAT, while paradoxically improving thermoregulation and promoting increased metabolic fitness in mice. These adaptations were mediated by the main effector of the ISR, ATF4, partially via the induction of FGF21 as a batokine. Nonetheless, the resistance to diet-induced obesity (DIO) and improvements in glucose homeostasis observed in OPA1-deficient mice occurred in an ATF4-dependent, but FGF21-independent manner [1]. Moreover, the molecular mechanisms leading to activation of the ISR remained incompletely understood. Therefore, in the present study we sought to investigate the molecular mechanisms mediating ISR induction and the mechanisms downstream of ATF4 promoting resistance to DIO in mice lacking OPA1 in thermogenic adipocytes.

The ISR can be induced by four different eIF2α kinases that act as early responders to disturbances in cellular homeostasis. Each kinase responds to distinct environmental and physiological stresses, which reflects their unique regulatory mechanisms [2]. We and others have shown induction of the ISR downstream of mitochondrial stress, but the mechanisms mediating this induction are incompletely understood [1, 6, 9, 10]. RNASeq data in OPA1-deficient BAT indicated an increase in ER stress pathways and activation of the UPR in OPA1 BAT KO. Therefore, here we tested the hypothesis that PERK, an eIF2α kinase shared by the UPR and the ISR, is required for ATF4 induction in OPA1-deficient BAT. Our data in mice demonstrated that, although ER stress is increased in response to OPA1 deletion, PERK is dispensable for eIF2α phosphorylation and ATF4 induction in OPA1-deficient BAT. Therefore, mice concomitantly lacking OPA1 and PERK in BAT display the same improvements in systemic metabolic health as observed in OPA1 BAT KO mice. These data suggest that an alternative ISR kinase is induced downstream of mitochondrial stress to activate the ISR in BAT when OPA1 is deleted. Indeed, a recent study reveals that mitochondrial dysfunction induces different paths to promote ISR activation, depending on both the nature of the mitochondrial stress and on the metabolic state of the cell [11]. Moreover, studies suggest that heme-regulated eIF2α kinase (HRI) is required to activate ATF4 in response to mitochondrial stress in HEK293T cells treated with oligomycin [12], and in embryonic and adult hearts of a mouse model of mitochondrial cardiomyopathy [13]. Whether the same mechanisms mediate ATF4 induction upon OPA1 deletion remain to be determined.

Transcriptome analysis also revealed that GDF15 was induced in OPA1-deficient BAT, which correlated with increased GDF15 serum levels. GDF15 is a member of the transforming growth factor-β superfamily. It acts through a recently identified receptor, an orphan member of the GFRα family called GFRAL, and signals through the Ret coreceptor [14]. Several studies have demonstrated that GDF15 mediates its effects on reducing food intake, body weight and adiposity largely by its actions on regions of the hindbrain [7, 15-17]. Indeed, systemic overexpression of GDF15 was shown to prevent obesity and insulin resistance by modulating metabolic activity and enhancing the expression of important thermogenic and lipolytic genes in BAT and WAT [18-20]. GDF15 is secreted by various organs such as liver [21], kidney [22] and skeletal muscle [6]. Recently, studies demonstrated that GDF15 can also be induced in brown adipocytes in response to thermogenic stimuli [23, 24] and prolonged high-fat feeding [25]. Like FGF21, GDF15 is also induced in response to mitochondrial stress downstream of the ISR [26]. Our data show that *Gdf15* mRNA levels were dramatically reduced in mice lacking both OPA1 and ATF4 in BAT, suggesting that ATF4 may indirectly regulate GDF15 induction in OPA1 BAT KO mice, perhaps via its downstream target, CHOP, which can bind to the GDF15 promoter [27]. Therefore, we hypothesized that GDF15 is downstream of ATF4 and required for the resistance to DIO observed in mice lacking OPA1 in BAT.

Our data in mice lacking both OPA1 and GDF15 in BAT revealed that GDF15 partially mediates the resistance to DIO, independently of changes in food intake. Our data suggest that BAT-derived GDF15 increases metabolic rates in OPA1 BAT KO mice fed obesogenic diets, thereby attenuating weight gain. Interestingly, although GDF15 effects on body weight were only partial, the improvements in glucose disposal and hepatic steatosis observed in OPA1 BAT KO mice were completely abrogated in the absence of BAT GDF15. Paradoxically, insulin sensitivity remained improved. In the context of hepatic mitochondrial dysfunction, which was associated with increased FGF21 and GDF15 in the liver and in the circulation, GDF15 was shown to regulate changes in body and fat mass and to protect against hepatic steatosis in DIO but had no effect on glucose disposal. Instead, FGF21 was required to increase insulin sensitivity, energy expenditure and UCP1-mediated thermogenesis in inguinal adipose tissue (iWAT) in regular chow-fed mice [5], which is similar to what we previously reported for OPA1 BAT KO mice [1]. FGF21 and GDF15 induction and secretion were also reported in a model of mitochondrial stress in adipose tissue (both BAT and WAT). In this study, under regular chow-fed conditions, neither GDF15 nor FGF21 appeared to regulate whole-body metabolism.

Conversely, similar to our study, long-term induction of GDF15 attenuated progression of obesity by increasing energy expenditure in DIO, while FGF21 was dispensable for this phenotype [26]. Together, our study suggests FGF21 and GDF15 play fundamentally distinct roles in regulating energy metabolism and glucose homeostasis in different metabolic states. Our data also support a predominant role for GDF15 rather than FGF21 in promoting resistance to DIO in models of tissue-specific mitochondrial stress. Some discrepancies between our model and similar models in the literature may stem from differences in the nature of the mitochondrial stress, and the fact that we deleted FGF21 [1] and GDF15 in a tissue-specific manner, rather than globally. Also, additional methodological details such as age, and duration of feeding might have contributed to differences in our overall conclusions. Because GDF15 is induced in BAT in response to prolonged high-fat feeding [25], it is possible that BAT-derived GDF15 may regulate energy metabolism and glucose homeostasis during overfeeding. Future studies in mice with selective deletion of GDF15 alone in thermogenic adipocytes are required to determine its role in BAT-mediated systemic metabolic adaptations to DIO and cold exposure.

GDF15 has been shown to be induced in brown adipocytes in response to thermogenic stimuli, although the role for GDF15 in thermogenesis in unclear [23, 24]. Because OPA1 BAT KO mice have improved thermoregulation, we tested the role of BAT-derived GDF15 for adaptive thermogenesis. The increase in baseline core body temperature observed in OPA1 BAT KO mice was still observed when GDF15 was deleted, suggesting FGF21 might play a predominant role on thermoregulation under ambient temperature conditions [1]. However, in response to cold, mice lacking both OPA1 and GDF15 became severely hypothermic as early as 4 hours into the cold exposure (4 °C) protocol. Surprisingly, these mice had increased markers of browning in white adipose tissue (WAT) after 5 hours of cold exposure but lacked the increase in sympathetic activation observed in OPA1 BAT KO mice. Our data suggest that, at least in the context of mitochondrial stress, GDF15 expression in thermogenic adipocytes plays a critical role in maintaining core body temperature in cold-exposed mice, likely by regulating sympathetic activation of sub-cutaneous white adipose tissue. Our results also demonstrate that both, increased browning and increased sympathetic input are required for proper thermoregulation during cold exposure in mice with impaired BAT thermogenesis. Future studies will be required to determine if the requirement for GDF15 to increase sympathetic activation of iWAT is mediated by central mechanisms.

In conclusion, our study has ruled out a role for PERK in inducing the ISR in response to OPA1 deletion in BAT. Furthermore, we also unveiled a role for GDF15 in attenuating DIO and improving glucose clearance and hepatic steatosis in OPA1 BAT KO mice. To our knowledge, this is the first demonstration that BAT-derived GDF15 plays a role on energy metabolism, glucose homeostasis and thermoregulation in the context of mitochondrial stress in BAT. Our data also shows that GDF15 may exert its effects independently of changes in food intake but predominantly by increasing energy expenditure. The specific molecular mechanisms downstream of GDF15 regulating these metabolic adaptations remain to be elucidated.

## Material and Methods

### Key Resources Table

| Key Resources Table |  |  |  |  |
| --- | --- | --- | --- | --- |
| Reagent type (species) or resource | Designation | Source or reference | Identifiers | Additional information |
| strain, strain background (Mouse, C57Bl/6J) | Murine Models | Jackson Laboratories | JAX Stock #025124 | Tag (Ucp1-cre)<sup>1</sup>Evd<sup>r</sup>; male and female |
| strain, strain background (Mouse, C57Bl/6J) | Murine Models | Jackson Laboratories | JAX Stock #023066 | *Eif2ak3*<sup>tm1.2Drc</sup>/J; male and female |
| antibody | anti-OPA1 (Mouse polyclonal) | BD Biosciences | #612606 | WB 1:1000, primary |
| antibody | Anti-GAPDH | Cell Signaling Technology | #2118 | WB 1:1000, primary |
| antibody | anti-UCP1 (Mouse monoclonal) | Abcam | #Ab10983 | WB 1:1000, primary<br>Histology 1:250 |
| antibody | Anti-α-tubulin | Sigma | #T9026 | WB 1:1000, primary |
| antibody | anti-β-actin | Sigma | #A2066 | WB 1:1000,<br>primary |
| antibody | Anti-phosphorylated eIF2α serine 51 | Cell Signaling Technology | 3597 | WB 1:1000,<br>primary |
| antibody | anti-eIF2α | Santa Cruz Biotechnology | #SC81261 | WB 1:1000,<br>primary |
| antibody | Anti-PERK | Cell Signaling Technology | C33E10 | WB 1:1000,<br>primary |
| antibody | IRDye 800CW anti-mouse | LI-COR | #925-32212 | WB<br>1:10,000,<br>secondary |
| antibody | Alexa Fluor anti-rabbit 680 | Invitrogen | #A27042 | WB<br>1:10,000,<br>secondary |
| commercial assay or kit | RNeasy kit | Qiagen Inc | #74104 | commercial assay or kit |
| commercial assay or kit | EnzyChrom™ Triglyceride Assay Kit | BioAssay Systems | #ETGA-200 |  |
| commercial assay or kit | Mouse/Rat Fibroblast Growth Factor 21 ELISA | Biovendor | #RD291108200R |  |
| commercial assay or kit | Quantikine GDF15 ELISA | R&D Systems | #MGD150 |  |
| commercial assay or kit | Ultra-Sensitive Mouse Insulin ELISA Kit | Chrystal Chem | #90080 |  |
| commercial assay or kit | High-Capacity cDNA reverse Transcription Kit | Applied Biosystems | #4368814 |  |
| software, algorithm | Graphpad Prism Software | Graphpad Software, La Jolla, CA, USA | Version 8.0.0 for Windows |  |
| other | Chow, standard | Harlan Teklad | 2029X |  |
| other | Chow, 60% HFD | Research Diets | D12492 |  |

### Appendix A - Primers

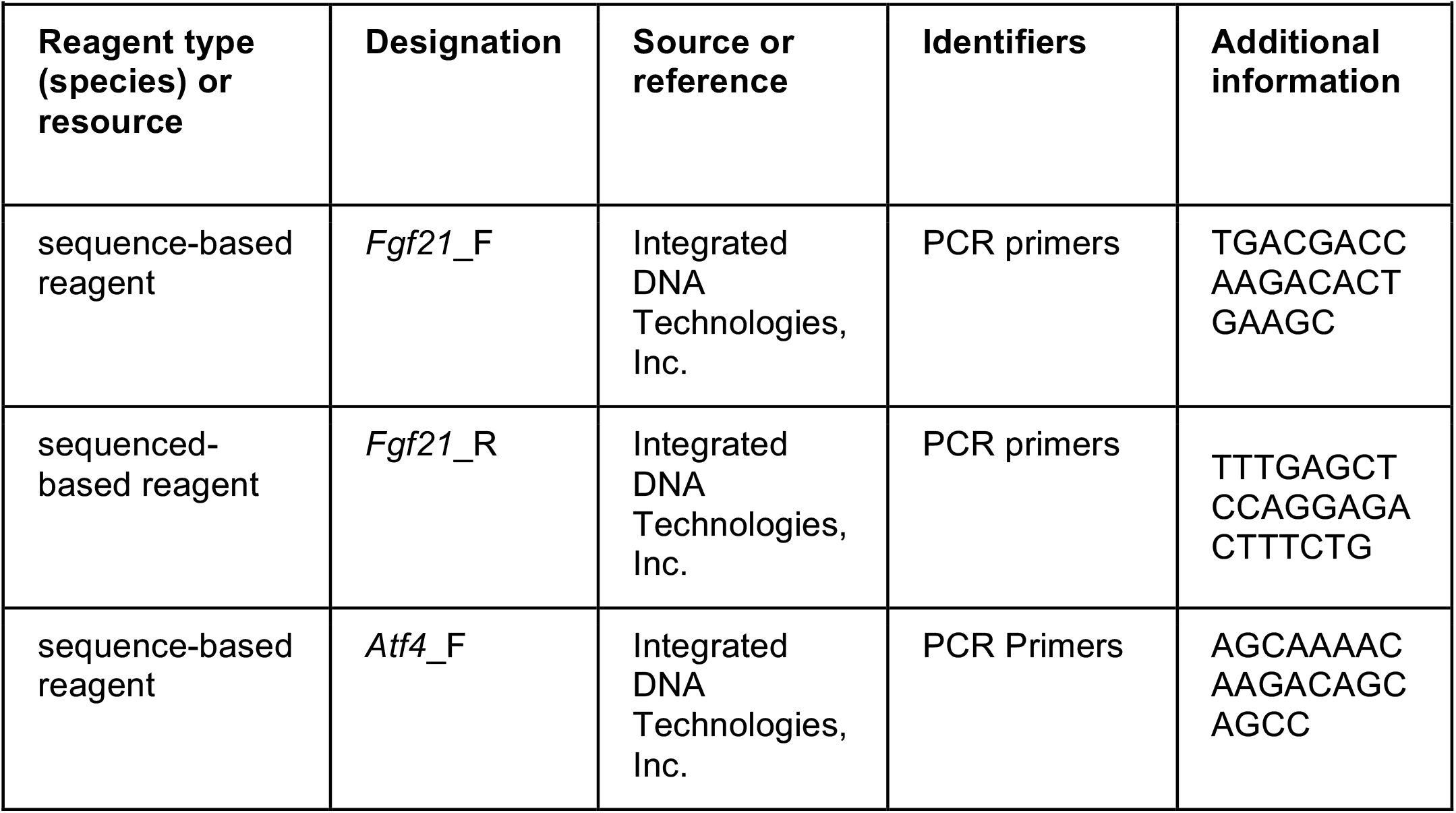
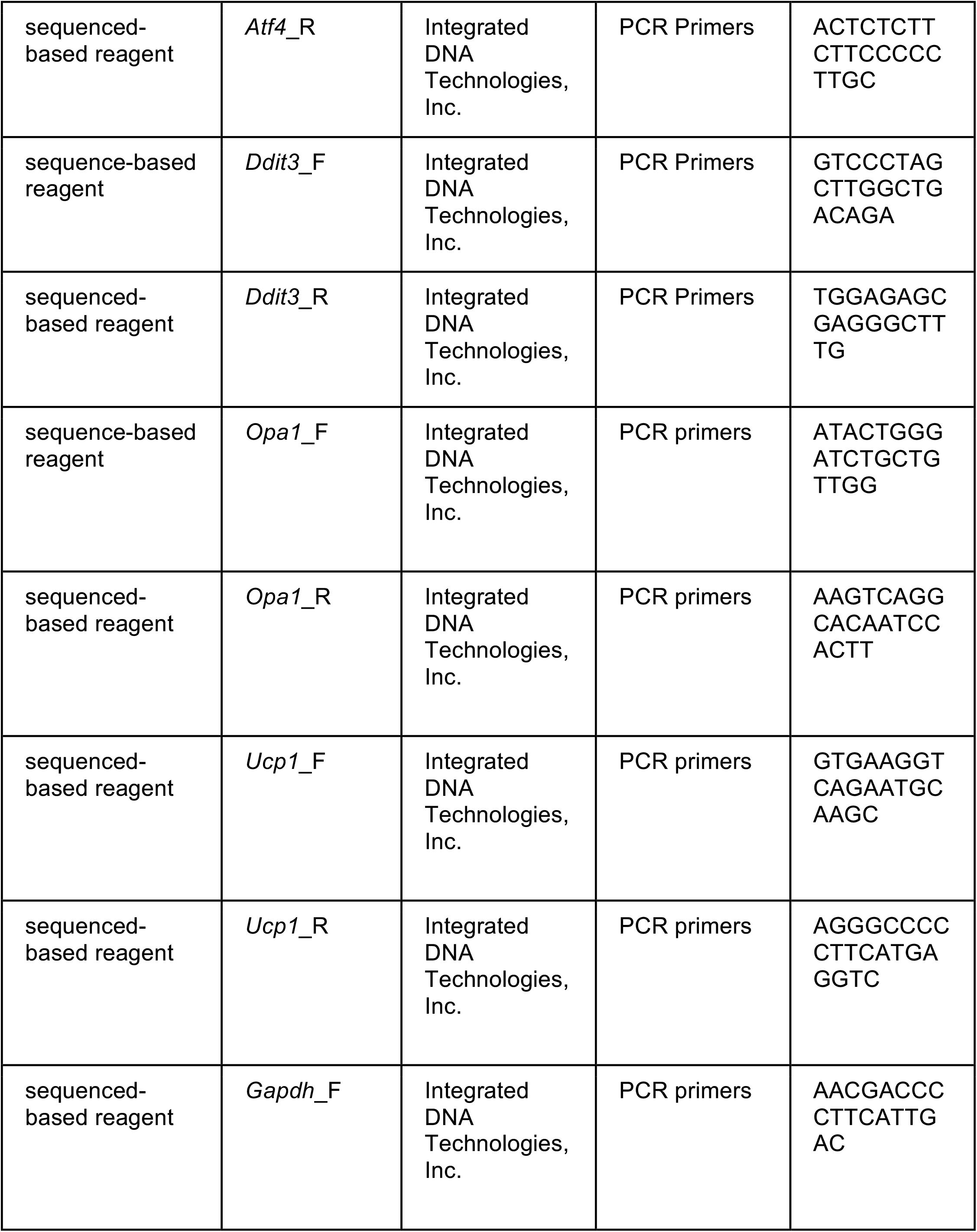
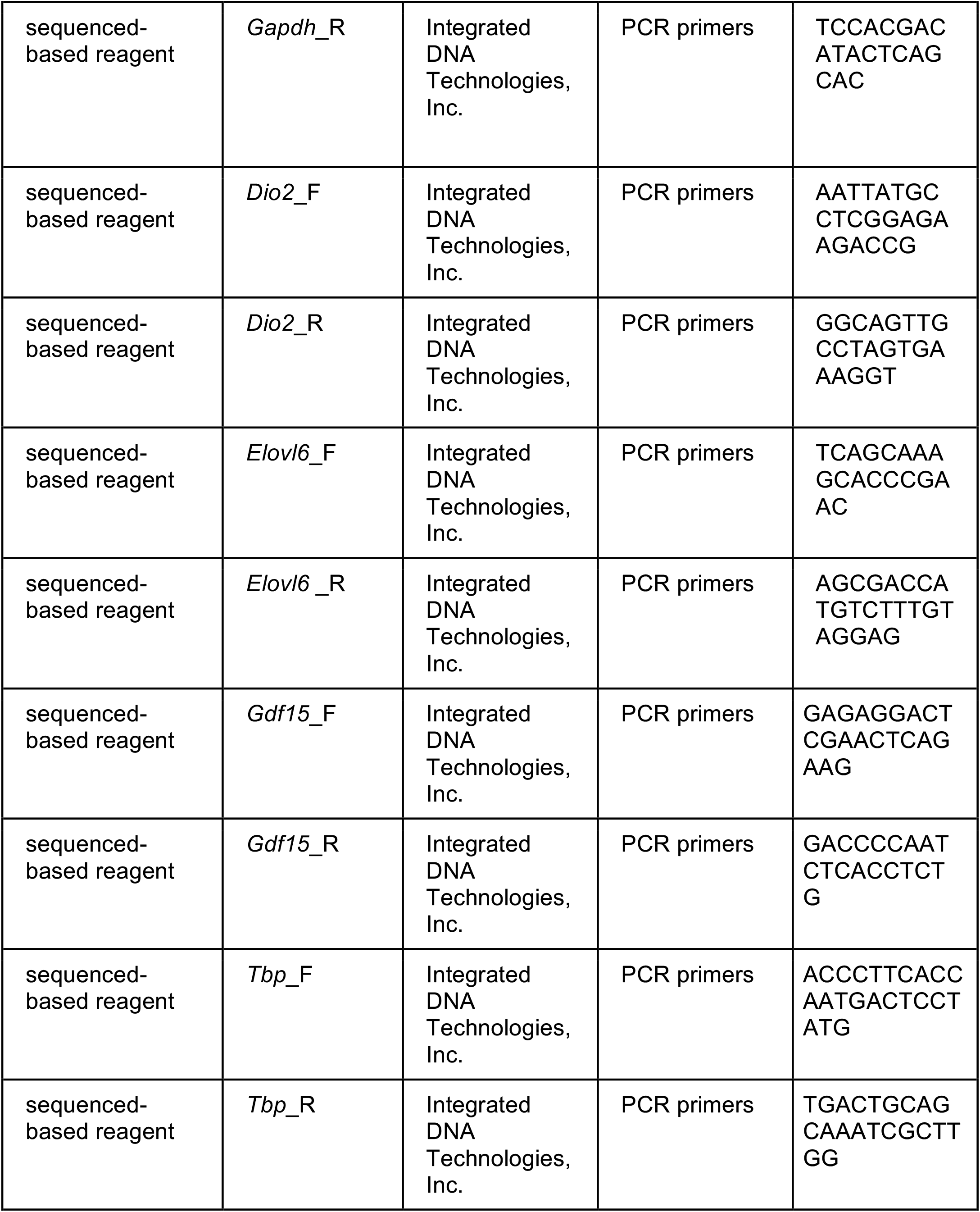

### Mouse Models

Experiments were performed in male and/or female mice on a C57Bl/6J background. OPA1^fl/fl^ mice [28], FGF21^fl/fl^ mice [29] and ATF4^fl/fl^ mice [30] were generated as previously described. GDF15 ^fl/fl^ mice were generated by Dr. Randy Seeley in the C657Bl/6J background and contain loxP sites flanking exon 2 of *Gdf15* gene. Briefly, to generate GDF15 ^fl/fl^ mice, the University of Michigan Transgenic Core injected Cas9 protein (Sigma-Aldrich, St. Louis, MO, USA), editing templates containing LoxP sites (IDT Ltd., Coralville, Iowa, USA) and two sgRNAs (Synthego, Redwood City, CA, USA) recognizing two sites upstream and downstream of exon two of *Gdf15* gene (target sequence: uuggauucacacaacccuag and aggaaaagggacauacagag) into the pro-nucleus of fertilized mouse embryos. Embryos were then implanted into pseudo pregnant dams. Resultant pups were screened for the presence of LoxP sites within the *Gdf15* genomic sequence. Surrounding sequences were then amplified and subjected to DNA sequencing.

Positive animals were bred to C57Bl6/J mice and the resultant pups were rescreened and resequenced prior to propagation. Subsequent animals were screened for *Gdf15* Flox by PCR (5’ forward: agccagagtaggacggatga; 5’ reverse: caattctgcttcaacccccg; 3’ forward: tgagcccttgggaggtagag; 3’ reverse: ggccacaaaccactctacga). Transgenic mice expressing cre recombinase under the control of the *Ucp1* promoter (Tg (Ucp1-cre)1Evdr) [31], and PERK ^fl/fl^ mice (*Eif2ak3*^*tm1*.*2Drc*^/J) [32] were acquired from the Jackson Laboratories (#024670 and #023066, respectively). Compound mutants were generated by crossing OPA1^fl/fl^ harboring the *Ucp1* Cre with FGF21^fl/fl^, ATF4^fl/fl^, PERK ^fl/fl^ or GDF15 ^fl/fl^ mice. Wild type (WT) controls for each compound mutant were mice harboring the respective homozygous floxed alleles but lacking the *Ucp1* Cre. Mice were weaned at 3 weeks of age and were kept on standard chow (2920X Harlan Teklad, Indianapolis, IN, USA). For diet-induced obesity studies, 6-week-old mice were fed a high-fat diet group (HFD; 60% Kcal from fat—Research Diets D12492) for 12 weeks. After 11 weeks of high-fed feeding, a subset of mice was placed in the Promethion System (Sable Systems International, Las Vegas, NV, USA) to measure changes in energy metabolism, food intake and locomotor activity. For the cold exposure experiments, mice were acclimated to 30 °C (thermoneutral temperature for mice) for 7 days prior to being cold-exposed. Unless otherwise noted, animals were housed at 22 °C with a 12-h light, 12-h dark cycle with free access to water and standard chow or special diets. All mouse experiments presented in this study were conducted in accordance with the animal research guidelines from NIH and were approved by the University of Iowa IACUC.

### Cold exposure experiments

For the 3-day cold exposure experiments, core body temperature was measured by telemetry (Respironics, G2 E-Mitter, Murrysville, PA, USA), as previously described [1]. For the acute cold exposure experiments, 12-week-old mice were initially individually housed in the rodent environmental chamber at 30 °C for 7 days. The initial temperature (t0) was recorded using a rectal probe (Fisher Scientific, Lenexa, KS, USA) at 7 am on day 8, after which the temperature was switched to 4 °C. Once the desired temperature was reached, we recorded rectal temperatures hourly for up to 5 hours of cold exposure.

### Glucose and insulin tolerance tests, nuclear magnetic resonance, and serum analysis

Glucose tolerance tests (GTT), insulin tolerance tests (ITT), and measurements of serum insulin and FGF21 levels were performed as previously described by us [1]. Serum GDF15 was measured using commercially available kits according to the manufacturers’ directions (R&D Systems, Minneapolis, MN, USA). Body composition was determined by nuclear magnetic resonance (NMR) in the Bruker Minispec NF-50 instrument (Bruker, Billerica, MA, USA).

### Analysis of triglyceride levels

Triglycerides levels were measured in liver of mice fed HFD for 12 weeks, using the EnzyChrom™ Triglyceride Assay Kit (BioAssay Systems, Hayward, CA, USA), as previously described [1].

### RNA extraction and quantitative RT–PCR

Total RNA was extracted from tissues with TRIzol reagent (Invitrogen) and purified with the RNeasy kit (Qiagen Inc, Germantown, MD, USA). Quantitative RT-PCR was performed as previously described [1]. Data were normalized to *Gapdh* or *Tbp* expression and results are shown as relative mRNA levels. qPCR primers were designed using Primer-Blast or previously published sequences [33].

### RNA Sequencing

RNA sequencing was performed in BAT of 7-week-old female and male OPA1 BAT KO mice by the Iowa Institute of Human Genetics: Genomics Division at the University of Iowa. Sequencing libraries were prepped using the Illumina TruSeq mRNA Stranded kit and sequenced on a HiSeq4000. Two approaches were used for read alignment, mapping, and quantification. First, a workflow using HISAT2 (v2.1.0), featureCounts (v1.6.3), and DESeq2 (v1.22.2) was performed [34-36]. The second approach used pseudo-alignment and quantification with Kallisto (v0.45.0) and DESeq2 for differential expression analysis [37]. Ingenuity® Pathway Analysis (IPA®) software from Qiagen was utilized for identification of potentially modified pathways.

Data visualization was performed using pheatmap and ggplot2 packages in R. RNA-seq data have been deposited to the GEO database under the accession number GSE218907.

### Western blot analysis

Immunoblotting analysis was performed in BAT and iWAT, as previously described [1]. Membranes were incubated with primary antibodies overnight and with secondary antibodies for 1 h, at room temperature. Fluorescence was quantified using the LiCor Odyssey imager.

## Data analysis

Unless otherwise noted, all data are reported as mean ± SEM. To determine statistical differences, Student’s *t*-test was performed for comparison of two groups, and Two-Way ANOVA followed by Tukey multiple comparison test was utilized when more than three groups were compared. A probability value of *P* ≤ 0.05 was considered significantly different. Statistical calculations were performed using the GraphPad Prism software (La Jolla, CA, USA). The association between oxygen consumption or energy expenditure and body mass were calculated by ANCOVA, using the CalR software [38]. The significance test for the “group effect” determined whether the two groups of interest were significantly different for the metabolic variable selected.

## Acknowledgment

This work was supported by AHA Scientist Development Grant 15SDG25710438 and NIH DK125405 to R.O.P.; by the Diabetes Research Training Program funded by the NIH (T32DK112751-01) to S.H.B; and by the NIH 1R25GM116686 to L.M.G.P. Metabolic phenotyping was performed at the Metabolic Phenotyping Core at the Fraternal Order of Eagles Diabetes Research Center. RNASeq was performed at the Genomics Division of The Iowa Institute of Human Genetics. We thank Dr. Matthew J. Potthoff and Dr. Christopher Adams for graciously sharing the FGF21^fl/fl^ mice and the ATF4^fl/fl^ mice, respectively.

## Author Contributions

R.O.P. conceived the project, wrote the manuscript, and coordinated all aspects of this work. J.J. and L.M.G.P. conducted the experiments, analyzed data, prepared figures and helped write the manuscript. E.T.W. analyzed the RNASeq data and helped prepare figures for the manuscript. A.M., S.H.B., K.K., J.K., J.C. aided with mouse phenotyping, processed tissues, and performed biochemical analysis. R.J.S. and E.D.A. provided essential materials and critical expertise. R.O.P., E.D.A, J.J., L.M.G.P. edited the manuscript.

## References

1. Pereira, R.O., et al., OPA1 deletion in brown adipose tissue improves thermoregulation and systemic metabolism via FGF21. Elife, 2021. 10.

2. Pakos-Zebrucka, K., et al., The integrated stress response. EMBO Rep, 2016. 17(10): p. 1374–1395.

3. Bao, X.R., et al., Mitochondrial dysfunction remodels one-carbon metabolism in human cells. Elife, 2016. 5.

4. Quiros, P.M., et al., Multi-omics analysis identifies ATF4 as a key regulator of the mitochondrial stress response in mammals. J Cell Biol, 2017. 216(7): p. 2027–2045.

5. Kang, S.G., et al., Differential roles of GDF15 and FGF21 in systemic metabolic adaptation to the mitochondrial integrated stress response. iScience, 2021. 24(3): p. 102181.

6. Ost, M., et al., Muscle-derived GDF15 drives diurnal anorexia and systemic metabolic remodeling during mitochondrial stress. EMBO Rep, 2020. 21(3): p. e48804.

7. Tsai, V.W.W., et al., The MIC-1/GDF15-GFRAL Pathway in Energy Homeostasis: Implications for Obesity, Cachexia, and Other Associated Diseases. Cell Metab, 2018. 28(3): p. 353–368.

8. Tsai, V.W., et al., Treatment with the TGF-b superfamily cytokine MIC-1/GDF15 reduces the adiposity and corrects the metabolic dysfunction of mice with diet-induced obesity. Int J Obes (Lond), 2018. 42(3): p. 561–571.

9. Forsstrom, S., et al., Fibroblast Growth Factor 21 Drives Dynamics of Local and Systemic Stress Responses in Mitochondrial Myopathy with mtDNA Deletions. Cell Metab, 2019. 30(6): p. 1040–1054 e7.

10. Ost, M., et al., Muscle mitochondrial stress adaptation operates independently of endogenous FGF21 action. Mol Metab, 2016. 5(2): p. 79–90.

11. Mick, E., et al., Distinct mitochondrial defects trigger the integrated stress response depending on the metabolic state of the cell. Elife, 2020. 9.

12. Guo, X., et al., Mitochondrial stress is relayed to the cytosol by an OMA1-DELE1-HRI pathway. Nature, 2020. 579(7799): p. 427–432.

13. Zhu, S., et al., Mitochondrial Stress Induces an HRI-eIF2alpha Pathway Protective for Cardiomyopathy. Circulation, 2022. 146(13): p. 1028–1031.

14. Breit, S.N., D.A. Brown, and V.W. Tsai, The GDF15-GFRAL Pathway in Health and Metabolic Disease: Friend or Foe? Annu Rev Physiol, 2021. 83: p. 127–151.

15. Xiong, Y., et al., Long-acting MIC-1/GDF15 molecules to treat obesity: Evidence from mice to monkeys. Sci Transl Med, 2017. 9(412).

16. Tsai, V.W., et al., GDF15 mediates adiposity resistance through actions on GFRAL neurons in the hindbrain AP/NTS. Int J Obes (Lond), 2019.

17. Tran, T., et al., GDF15 deficiency promotes high fat diet-induced obesity in mice. PLoS One, 2018. 13(8): p. e0201584.

18. Macia, L., et al., Macrophage inhibitory cytokine 1 (MIC-1/GDF15) decreases food intake, body weight and improves glucose tolerance in mice on normal & obesogenic diets. PLoS One, 2012. 7(4): p. e34868.

19. Chrysovergis, K., et al., NAG-1/GDF-15 prevents obesity by increasing thermogenesis, lipolysis and oxidative metabolism. Int J Obes (Lond), 2014. 38(12): p. 1555–64.

20. Kim, J.M., et al., NAG-1/GDF15 transgenic mouse has less white adipose tissue and a reduced inflammatory response. Mediators Inflamm, 2013. 2013: p. 641851.

21. Patel, S., et al., Combined genetic deletion of GDF15 and FGF21 has modest effects on body weight, hepatic steatosis and insulin resistance in high fat fed mice. Mol Metab, 2022. 65: p. 101589.

22. Baek, S.J. and T. Eling, Growth differentiation factor 15 (GDF15): A survival protein with therapeutic potential in metabolic diseases. Pharmacol Ther, 2019. 198: p. 46–58.

23. Flicker, D., et al., Exploring the In Vivo Role of the Mitochondrial Calcium Uniporter in Brown Fat Bioenergetics. Cell Rep, 2019. 27(5): p. 1364–1375 e5.

24. Campderros, L., et al., Brown Adipocytes Secrete GDF15 in Response to Thermogenic Activation. Obesity (Silver Spring), 2019.

25. Patel, S., et al., GDF15 Provides an Endocrine Signal of Nutritional Stress in Mice and Humans. Cell Metab, 2019. 29(3): p. 707–718 e8.

26. Choi, M.J., et al., An adipocyte-specific defect in oxidative phosphorylation increases systemic energy expenditure and protects against diet-induced obesity in mouse models. Diabetologia, 2020. 63(4): p. 837–852.

27. Chung, H.K., et al., GDF15 deficiency exacerbates chronic alcohol- and carbon tetrachloride-induced liver injury. Sci Rep, 2017. 7(1): p. 17238.

28. Zhang, Z., et al., The dynamin-related GTPase Opa1 is required for glucose-stimulated ATP production in pancreatic beta cells. Mol Biol Cell, 2011. 22(13): p. 2235–45.

29. Potthoff, M.J., et al., FGF21 induces PGC-1alpha and regulates carbohydrate and fatty acid metabolism during the adaptive starvation response. Proc Natl Acad Sci U S A, 2009. 106(26): p. 10853–8.

30. Ebert, S.M., et al., Stress-induced skeletal muscle Gadd45a expression reprograms myonuclei and causes muscle atrophy. J Biol Chem, 2012. 287(33): p. 27290–301.

31. Kong, X., et al., IRF4 is a key thermogenic transcriptional partner of PGC-1alpha. Cell, 2014. 158(1): p. 69–83.

32. Zhang, P., et al., The PERK eukaryotic initiation factor 2 alpha kinase is required for the development of the skeletal system, postnatal growth, and the function and viability of the pancreas. Mol Cell Biol, 2002. 22(11): p. 3864–74.

33. Kim, K.H., et al., Autophagy deficiency leads to protection from obesity and insulin resistance by inducing Fgf21 as a mitokine. Nat Med, 2013. 19(1): p. 83–92.

34. Kim, D., et al., Graph-based genome alignment and genotyping with HISAT2 and HISAT-genotype. Nat Biotechnol, 2019. 37(8): p. 907–915.

35. Liao, Y., G.K. Smyth, and W. Shi, featureCounts: an efficient general purpose program for assigning sequence reads to genomic features. Bioinformatics, 2014. 30(7): p. 923–30.

36. Love, M.I., W. Huber, and S. Anders, Moderated estimation of fold change and dispersion for RNA-seq data with DESeq2. Genome Biol, 2014. 15(12): p. 550.

37. Bray, N.L., et al., Near-optimal probabilistic RNA-seq quantification. Nat Biotechnol, 2016. 34(5): p. 525–7.

38. Mina, A.I., et al., CalR: A Web-Based Analysis Tool for Indirect Calorimetry Experiments. Cell Metab, 2018. 28(4): p. 656–666 e1.

